# Colon Cancer and SARS-CoV-2: Impact of ACE2 Expression in Susceptibility to COVID-19

**DOI:** 10.1101/2020.06.11.146878

**Authors:** Mohsen Ahmadi, Negin Saffarzadeh, Mohammad Amin Habibi, Fatemeh Hajiesmaeili, Nima Rezaei

## Abstract

Novel coronavirus disease (COVID-19) pandemic has become a global health emergency. Severe acute respiratory syndrome coronavirus 2 (SARS-CoV-2) interacts with angiotensin-converting enzyme 2 (ACE2) to enter the cells and infects diverse human tissues. It has been reported that a few conditions, including cancer, predispose individuals to SARS-CoV-2 infection and severe form of COVID-19. These findings led us to evaluate the susceptibility of colon adenocarcinoma (COAD) patients to SARS-CoV-2 infection by investigation of ACE2 expression in their tumor tissues. The expression analysis revealed that both mRNA and protein levels of ACE2 had increased in colon cancer samples than normal group. Next, the prognosis analysis has indicated that the upregulation of ACE2 was not correlated with patient survival outcomes. Further assessment displayed the hypomethylation of the ACE2 gene promoter in COAD patients. Surprisingly, this methylation status has a strong negative correlation with ACE2 gene expression. The functional enrichment analysis of the genes that had similar expression patterns with ACE2 in colon cancer tissues demonstrated that they mainly enriched in Vitamin digestion and absorption, Sulfur relay system, and Fat digestion and absorption pathways. Finally, we found that ACE2 gene expression had a significant association with the immune cell infiltration levels in COAD patients. In conclusion, it has plausible that COAD patients are more likely to be infected with SARS-CoV-2 and experience severe injuries. Moreover, COVID-19 would bring unfavorable survival outcomes of patients with colon cancer by the way of immune cell infiltration linked process. The present study highlights the importance of preventive actions for COAD patients during the COVID-19 pandemic.

## Introduction

Since the first report of severe acute respiratory syndrome coronavirus 2 (SARS-CoV-2) in China in December 2019, coronavirus disease (COVID-19), has affected about six million confirmed cases around the world by the end of May 2020 (1-3). The wide range of COVID-19 symptoms, such as fever, cough, tiredness, headache, diarrhea, and cardiac injury, implying that SARS-CoV-2 infects not only the respiratory tract but also various human organs (2, 4–8). SARS-CoV-2 interacts with angiotensin-converting enzyme 2 (ACE2) receptors to start infection of the cells (9–12). Interestingly, a higher prevalence of COVID-19 has shown in patients with diabetes, hypertension, and cardiovascular diseases (13, 14). Moreover, the mRNA levels of the *ACE2* gene are particularly high in many human tissues including the lung, heart, liver, kidney, stomach, and colon (15–17). Albeit the exact underlying mechanism is unknown, the expression levels of ACE2 have introduced to play a role in the increased susceptibility of diabetic and hypertensive patients to SARS-CoV-2 infection. Therefore, the expression levels of ACE2 seem to have a prominent effect on the individuals’ risk for the COVID-19.

Colon adenocarcinoma (COAD), as the most common type of colon cancer, ranks fourth most common cancer by incidence and the second leading cause of cancer-related deaths worldwide (18, 19). Although the diagnosis and treatment of COAD patients are much improved, the survival rate of these patients remains unsatisfactory, mostly due to the late-diagnosis (20, 21). Increasing data has indicated the correlation of the immune cell infiltration in colon cancer with the patients’ clinical outcomes (22). It has been well-established that cancerous patients are more likely to be infected with SARS-CoV-2 than those without, because of anticancer treatments and altered immune state. Besides, they appear to have severe COVID-19 symptoms in comparison to the general population (23, 24). These findings led us to investigate the susceptibility of colon cancer patients to SARS-CoV-2 infection by analyzing the expression levels of the ACE2 gene in these patients as a fundamental component for the SARS-CoV-2 pathogenesis. Indeed, we tested the possibility of changing the prognosis of COAD patients through immune cell infiltration dependent mechanism after infection by COVID-19.

In the present study, we used several databases to investigate the gene expression, prognostic merit, and the DNA methylation status of the *ACE2* gene in colon adenocarcinoma. We also conducted an enrichment analysis of ACE2 co-expressed genes. Then, the association of the *ACE2* gene expression with immune cell infiltration level in COAD patients was examined.

## Materials and Methods

### Expression analysis

The expression levels of the human *ACE2* gene in the colon cancer and corresponding control tissues in the TCGA-COAD (Colon Adenocarcinoma) cohort, were assessed by the Gene Expression Profiling Interactive Analysis (GEPIA) 2 (http://gepia2.cancer-pku.cn/#analysis) database (25). Adjusted *p*-value <0.01, and log2 fold change | log2fc | >1 were set as cut of criterion. Besides, we applied the protein expression analysis for ACE2 in normal tissues and colon cancer, utilizing the data from Clinical Proteomic Tumor Analysis Consortium (CPTAC) Confirmatory/Discovery dataset through UALCAN (http://ualcan.path.uab.edu/analysis-prot.html) database (26).

### Prognosis analysis

We employed two online databases to perform the survival analysis for the *ACE2* gene in colon cancer patients. The OncoLnc database (http://www.oncolnc.org/) was utilized to conducting the Overall Survival (OS) analysis, by the expression level of the *ACE2* gene, based on the data of TCGA-COAD cohort (27). In addition, we evaluated the prognostic significance of *ACE2* gene expression, using the Meta-survival analysis option in the GENT2 database (http://gent2.appex.kr/gent2/) (28). The auto-select best cutoff was set for both databases.

### DNA methylation analysis

We investigated the DNA methylation status of the *ACE2* gene in the COAD cases of the TCGA-COAD cohort through the UALCAN database. Furthermore, the association between the gene expression and promoter methylation of the *ACE2* gene in the normal and primary colon cancer tissues, was explored by the DNMIVD database(http://119.3.41.228/dnmivd/) (29).

### Functional enrichment analysis

Based on the data of the TCGA-COAD cohort, we used the GEPIA 2 database to provide a group of genes that have a similar expression pattern with ACE2 in colon cancer. Then, the data of Gene Ontology (GO) and Kyoto Encyclopedia of Genes and Genomes (KEGG) pathway analysis for these co-expressed genes were obtained from the Enrichr (http://amp.pharm.mssm.edu/Enrichr) database (30). *p*-value less than 0.05 was set as a cut off criterion.

### Immune cell infiltration analysis

We utilized the CIBERSORT algorithm through TIMER 2.0 database (http://timer.cistrome.org/) (31) to evaluate the correlation of the *ACE2* expression with immune cells infiltration in COAD, including CD8+ T cell, activated Memory CD4+T cell, resting Memory CD4+T cell, Naive CD4+ T cell, Regulatory T cell (Tregs), Follicular helper T cell, Gamma delta T cell, Memory B cell, Naive B cell, Plasma B cell, Neutrophils, Monocytes, Eosinophils, M0 Macrophages, M1 Macrophages, M2 Macrophages, activated Myeloid Dendritic Cell (DC), resting Myeloid DC, activated Natural Killer (NK) cell, and resting NK cell. The correlation *p*-value adjusted by tumor purity.

## Results

### Expression analysis

First, we conducted the expression analysis using GEPIA 2 database. As depicted in Figure 1, pan-cancer analysis of the human ACE2 gene, revealed that mRNA levels of ACE2 increased in several tumors, especially in colon cancer (Figure 1A). Box plot analysis detected an overexpression level of ACE2 in the tissues of colon adenocarcinoma (Figure 1B, Adjusted P<0.01), compared with the normal tissues. The protein expression analysis results further showed that the expression level of ACE2 protein was higher in the cancer tissues than that in the controls (Figure 1C, P=7.53e-04).

**Figure 1.**
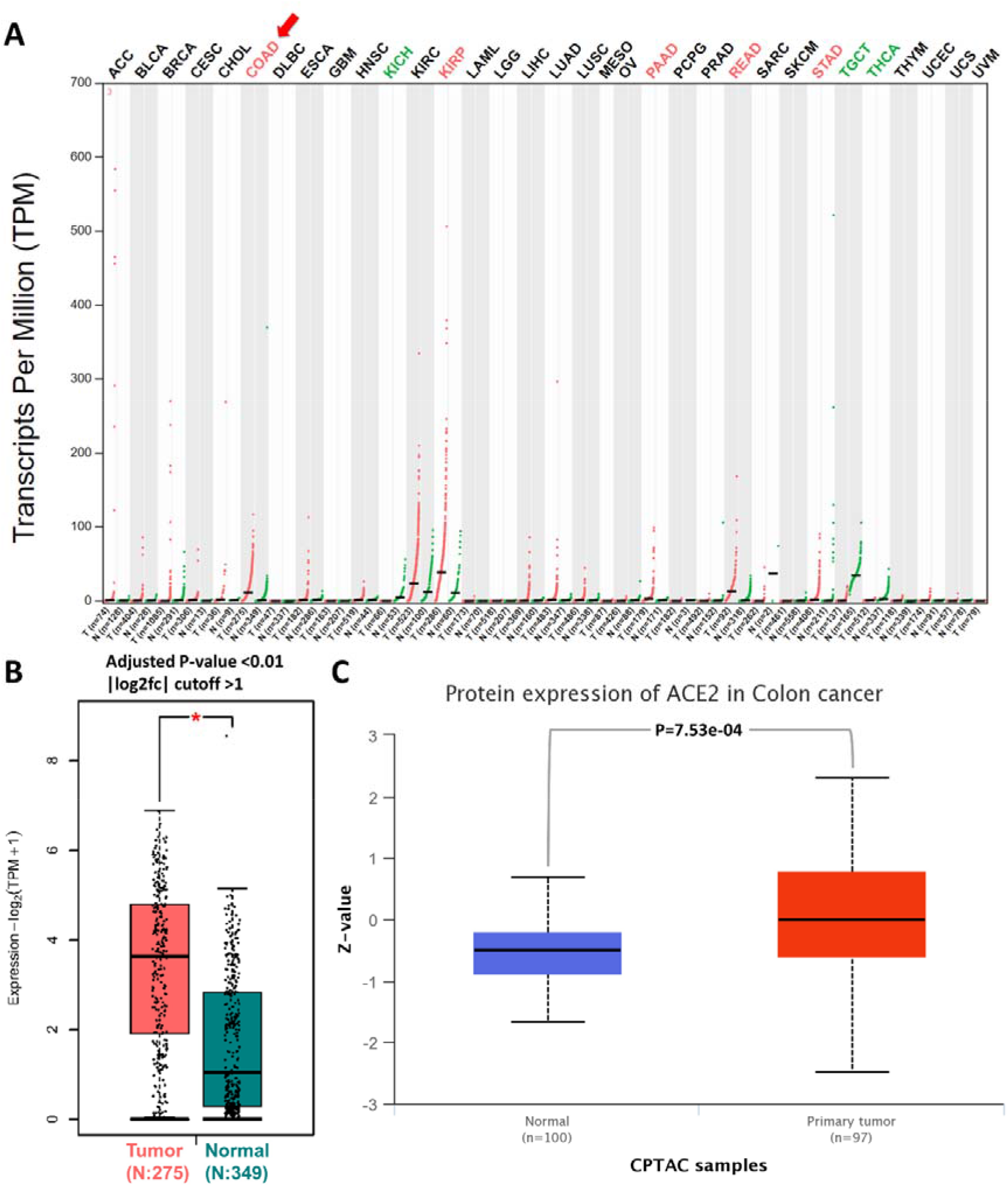
The expression analysis of the ACE2 in colon cancer. We demonstrated the expression level of *ACE2* gene in multiple cancers (A), especially in colon adenocarcinoma tissues and corresponding normal tissues (B), using GEPIA 2 database. The protein expression levels of ACE2 in normal tissue and colon cancer tissue were also obtained by UALCAN database (C).

### Prognosis analysis

We carried out a correlation analysis between the *ACE2* gene expression and the prognosis outcome of colon cancer patients based on the TCGA-COAD cohort data from OncoLnc database and survival data of multiple GEO datasets in GENT2 platform. The results demonstrated not a significant correlation between elevated expression of the ACE2 gene and overall survival of patients with colon cancer (Figure 2).

**Figure 2.**
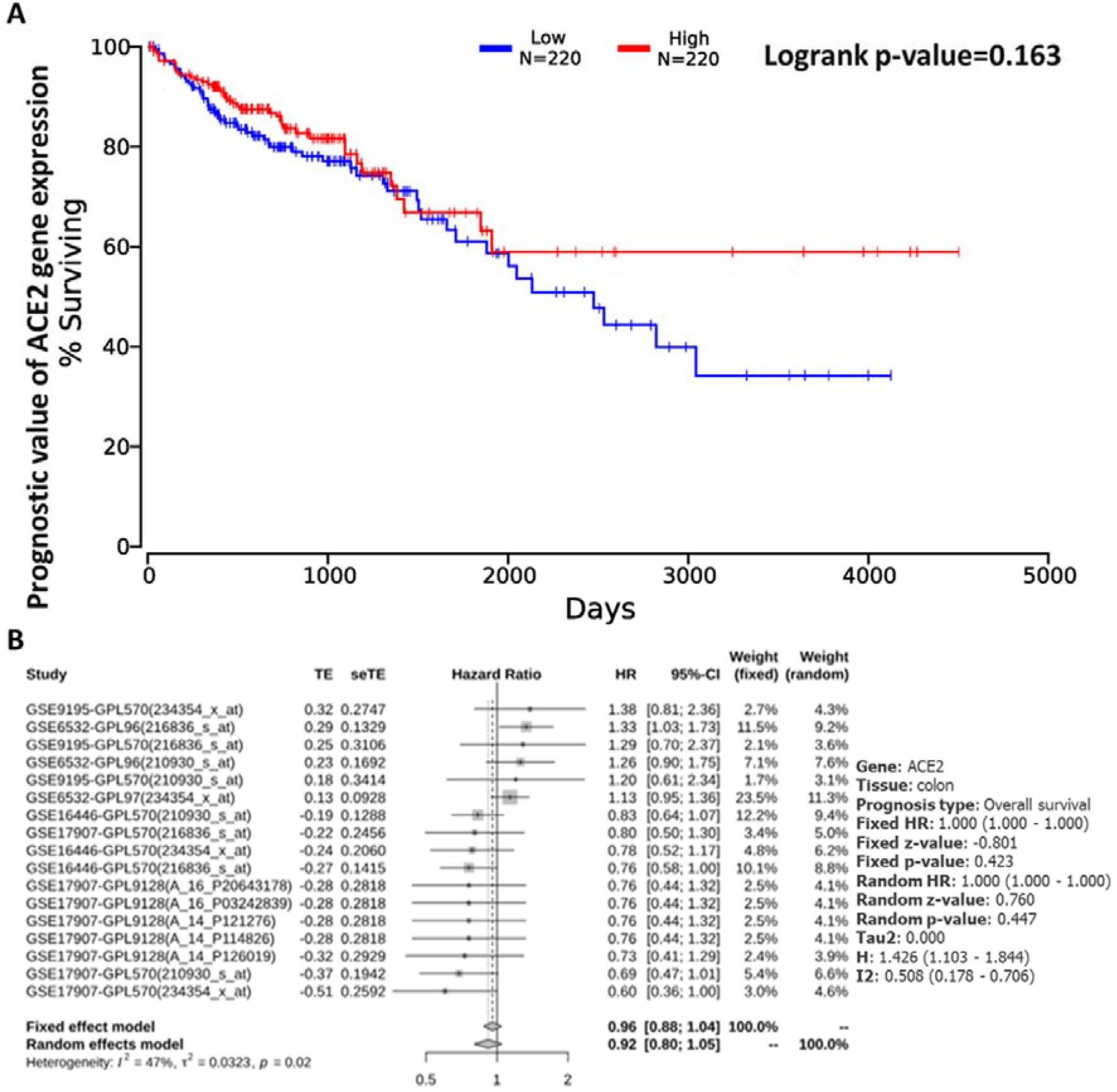
Survival analysis of the *ACE2* gene for colon cancer cases. We used the OnoLnc, and GENT2 databases to assess the overall survival analyses Based on the TCGA-COAD cohort data (A) and survival data of multiple GEO datasets (B), respectively.

### DNA methylation analysis data

We further studied the potential etiology of the elevated *ACE2* expression in colon cancer patients. So, we used UALCAN database to determine the methylation status of *ACE2* promoter in COAD. Interestingly, the methylation status of the *ACE2* promoter in COAD was significantly reduced, when compared with that in normal tissue (Figure 3A). Meanwhile, DNMIVD database analysis indicated that the promoter region of the *ACE2* gene was hypomethylated in COAD tumor tissues (Figure 3B). Notably, the Pearson and Spearman correlation analysis revealed a significant negative correlation between the expression of the *ACE2* mRNA and methylation status of *ACE2* gene in COAD (Figure 3C, 3D). Thus, DNA methylation may be involved in the molecular mechanism of the *ACE2* gene overexpression in tumor tissues and in the pathogenesis of colon cancer.

**Figure 3.**
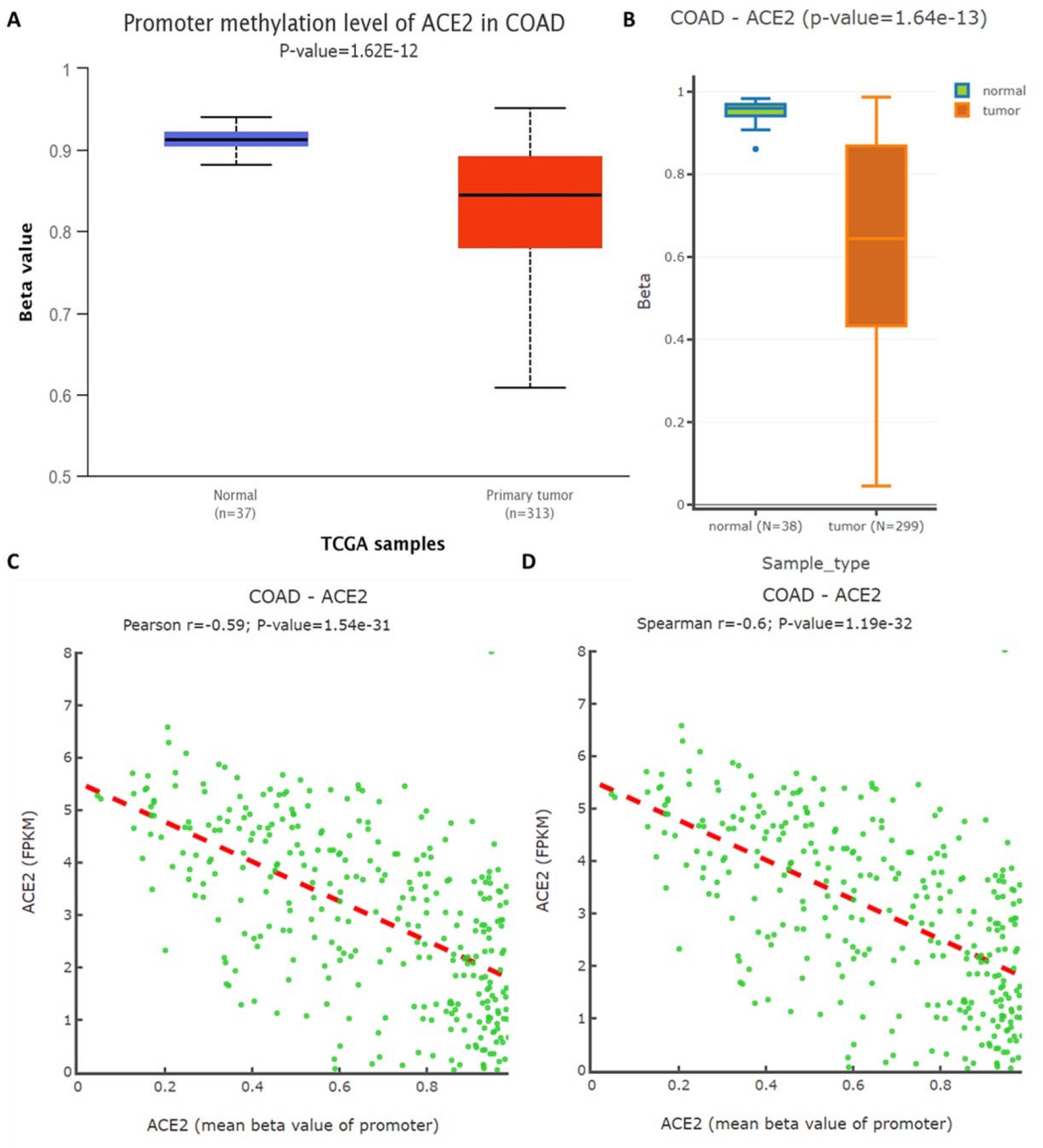
The promoter methylation status of *ACE2* gene in Colon Adenocarcinoma (COAD). DNA methylation analysis by UALCAN showed a lower methylation level for *ACE2* gene in COAD (A). We also detected the hypomethylation of *ACE2* promoter in COAD (B), and strong negative correlation between mRNA expression and methylation status of *ACE2* gene in COAD (C, D), using DNMIVD database.

### Functional analysis of co-expressed genes of ACE2 in colon cancer

The co-expression analysis using GEPIA 2 database provided a total of 100 associated genes for ACE2 in colon cancer (Table S1). The GO analysis data of Figure 4A, 4C identified the related biological process (vitamin transport, vitamin transmembrane transport, regulation of reactive oxygen species metabolic process), cellular component (cortical cytoskeleton, cytoskeleton, histone acetyltransferase complex), and molecular function (ribosomal protein S6 kinase activity, sulfurtransferase activity, solute:sodium symporter activity). KEGG pathway enrichment analysis data further exploited several enriched pathways, such as Vitamin digestion and absorption, Sulfur relay system, and Fat digestion and absorption pathways (Figure 4D).

**Figure 4.**
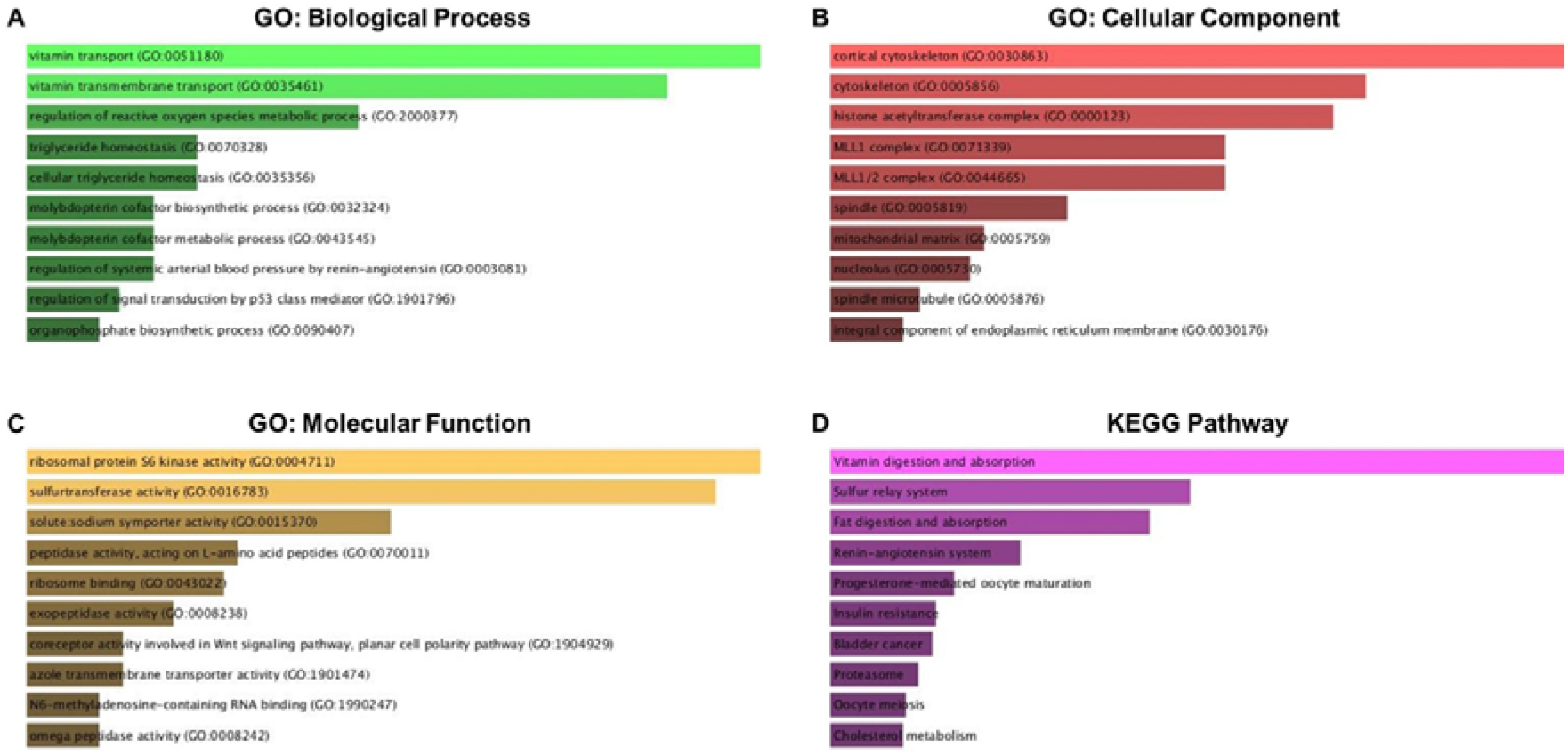
Data of GO and KEGG enrichment analysis. Based on the *ACE2*-correlated genes, the enriched information of biological process (A), cellular component (B), and molecular function (C) in GO analysis and KEGG pathway (D) were obtained using the Enrichr database. GO: gene ontology; KEGG: Kyoto encyclopedia of genes and genomes.

### Immune cell infiltration analysis data

We investigated whether the transcription levels of the *ACE2* gene play a role in the etiology of colon cancer through immune cell infiltration. As listed in Table 1, the expression of the *ACE* gene in colon cancer positively correlated with the infiltration level of the following immune cells: Memory B cell, Plasma B cell, activated myeloid DC, and resting myeloid DC. Moreover, the infiltration level of CD8+, T cell, and activated NK cells negatively correlated with the ACE2 gene expression levels (Figure S1).

**Table 1.** Correlation between *ACE2* gene expression and infiltration level of immune cells in COAD using TIMER 2.0 database.

| **Cell Type** | **Rho value** | **P-value** |
| --- | --- | --- |
| **CD8+ T cell** | **-0.1975** | **<0.0009** |
| activated Memory CD4+ T cell | 0.0025 | 0.9658 |
| resting Memory CD4+ T cell | 0.1079 | 0.0734 |
| Naive CD4+ T cell | 0.0355 | 0.5559 |
| Regulatory T cell (Tregs) | 0.0305 | 0.6136 |
| Follicular helper T cell | -0.02906 | 0.630746 |
| Gamma delta T cell | -0.10829 | 0.072473 |
| **Memory B cell** | **0.1300** | **<0.0308** |
| Naive B cell | -0.0261 | 0.6647 |
| **Plasma B cell** | **0.2301** | **<0.0001** |
| Neutrophils | -0.0319 | 0.5966 |
| Monocytes | 0.0715 | 0.2358 |
| Eosinophils | 0.085887 | 0.154732 |
| M0 Macrophages | 0.0450 | 0.4556 |
| M1 Macrophages | -0.0940 | 0.1191 |
| M2 Macrophages | -0.0724 | 0.2301 |
| **activated Myeloid Dendritic Cell (DC)** | **0.1859** | **<0.0019** |
| **resting Myeloid DC** | **0.1185** | **<0.0491** |
| **activated Natural Killer (NK) cell** | **-0.2443** | **<0.0000** |
| resting NK cell | 0.0854 | 0.1567 |

COAD: Colon Adenocarcinoma, rho value of Spearman's correlation, correlation p-value was adjusted by purity.

## Discussion

The COVID-19 pandemic is now widely spreading across the world and has become an international health emergency (32). The clinical manifestations of COVID-19 are varied, ranging from pulmonary-associated injuries to multi-organ failure, indicating that SARS-CoV-2 affects diverse human tissues (33). The SARS-CoV-2 binds to the ACE2 receptors to infect the host cells (6). It has reported that cancer patients have a higher risk of being infected by SARS-CoV-2 and developing severer symptoms, mainly because of compromised immune systems and the malignancy state (24). So, the ACE2 receptor maybe serves as an essential factor for the pathogenesis of COVID-19 (34). The present study explored the biological links of the human ACE2 gene expression with SARS-CoV-2 in colon cancer patients from different perspectives.

We first used the GEPIA 2 database and detected the overexpression of *ACE2* mRNA in primary tissues when compared with controls. The protein expression level analysis by the UALCAN database displayed that the protein levels of ACE2 were increased in COAD patients. We further explored the prognostic significance of the ACE2 gene expression in patients with colon adenocarcinoma utilizing the OnoLnc and GENT2 databases. The survival analysis showed no apparent association between the *ACE2* gene mRNA levels and the patient’s overall survival outcome. Subsequently, using co-expression analysis of GEPIA 2 database, we obtained the ACE2 co-expressed genes in colon cancer tissues, and then carried out a series of functional annotation and enrichment analyses. Our data demonstrated that the ACE2 mostly enriched in Vitamin digestion and absorption, Sulfur relay system, and Fat digestion and absorption pathways. Finally, we evaluated the correlation of *ACE2* gene expression with immune cell infiltration in patients with colon adenocarcinoma. The data retrieved from the TIMER database indicated a considerable correlation between the ACE2 gene expression and the infiltration level of CD8+ T cell, Memory B cells, Plasma B cells, activated myeloid DCs, resting myeloid DCs, and activated NK cells in patients with colon cancer.

The detected upregulation of the ACE2 gene in tumor tissues suggested that COAD patients were more vulnerable to SARS-CoV-2 infection. Several pieces of evidence highlighted the role of tumor-infiltrating immune cells in colon cancer patients’ survival (35, 36). For example, Wu D *et al*. uncovered the remarkable association of memory B cells with colon cancer survival risk (22). Surprisingly, our data unearthed a considerable correlation between ACE2 expression and immune cell infiltration in COAD. It has indicated that the SARS-CoV genome sequence has high homology with SARS-CoV-2 (37) Moreover, Kuba *et al*. showed the SARS-CoV decreased the expression of ACE2 in the lung cells (38). These findings unveiled that the levels of ACE2 in colon tumor tissues similarly be reduced after COVID-19 infection. If so, the prognosis of colon cancer patients may be influenced by SARS-CoV-2 infection, since we found a positive association between ACE2 expression and Immune cell infiltration in colon cancer patients.

## Conclusions

All data together, the ACE2 levels had increased in colon cancer tissues. Hence, these patients appear to be more susceptible to COVID-19. Furthermore, the SARS-CoV-2 infection presumably reduced the ACE2 level in tumor tissues and altered immune cell infiltration level, which would bring poor prognosis for patients with colon cancer. Our findings highlight the value of protective actions for COAD patients during the COVID-19 pandemic era.

## Acknowledgement

This study was supported by a grant from the Booali USERN Office, Booali Medical Diagnostic Laboratory.

## Authors’ contributions

NR supervised the project. MA has contribution in conceptualization, formal analysis, methodology, and visualization. NS did project administration and validation. FH did the formal analysis. MA and MAH drafted the manuscript. NR, NS, and FH critically revised it. All the authors approved the final draft of paper.

## Abbreviations

ACE2: Angiotensin-Converting Enzyme 2
COAD: Colon Adenocarcinoma
COVID-19: Coronavirus Disease of 2019
CPTAC: Clinical Proteomic Tumor Analysis Consortium
DC: Dendritic Cell
DNMIVD: DNA Methylation Interactive Visualization Database
GENT2: Gene Expression database of Normal and Tumor tissues 2
GEPIA: Gene Expression Profiling Interactive Analysis
GO: Gene Ontology
KEGG: Kyoto Encyclopedia of Genes and Genomes
NK cell: Natural Killer cell
OS: Overall Survival
SARS-CoV: Severe Acute Respiratory Syndrome Coronavirus
SARS-CoV-2: Severe Acute Respiratory Syndrome Coronavirus 2
TCGA: The Cancer Genome Atlas
Treg: Regulatory T cell

