## Supplementary material for "Colon Cancer and SARS-CoV-2: Impact of ACE2 Expression in Susceptibility to COVID-19": Figure S1

Table S1. The top 100 co-expressed genes of ACE2 in colon adenocarcinoma, obtained from the GEPIA 2 database.

| Gene Symbol | Gene ID | PCC |
| --- | --- | --- |
| KB-1732A1.1 | ENSG00000253669.3 | 0.58 |
| PLA2G12B | ENSG00000138308.5 | 0.58 |
| CEBPA | ENSG00000245848.2 | 0.57 |
| OIT3 | ENSG00000138315.12 | 0.56 |
| RPS6KA6 | ENSG00000072133.10 | 0.56 |
| SRPX2 | ENSG00000102359.5 | 0.55 |
| TP53RK | ENSG00000172315.5 | 0.54 |
| CSE1L | ENSG00000124207.16 | 0.54 |
| PLAGL2 | ENSG00000126003.6 | 0.53 |
| CASK | ENSG00000147044.20 | 0.53 |
| MOCS3 | ENSG00000124217.4 | 0.53 |
| TSPAN6 | ENSG00000000003.14 | 0.53 |
| RPS6KA3 | ENSG00000177189.12 | 0.53 |
| POFUT1 | ENSG00000101346.11 | 0.52 |
| ABCC6P2 | ENSG00000255277.3 | 0.52 |
| ID1 | ENSG00000125968.8 | 0.52 |
| RP11-150O12.3 | ENSG00000254290.1 | 0.52 |
| PRR15 | ENSG00000176532.3 | 0.52 |
| SLC2A8 | ENSG00000136856.17 | 0.52 |
| PXMP4 | ENSG00000101417.11 | 0.51 |
| NFS1 | ENSG00000244005.12 | 0.51 |
| C20orf24 | ENSG00000101084.16 | 0.51 |
| PSMA7 | ENSG00000101182.14 | 0.51 |
| AC106876.2 | ENSG00000222001.2 | 0.51 |
| EIF2S2 | ENSG00000125977.6 | 0.51 |
| GGH | ENSG00000137563.11 | 0.5 |
| ZSWIM3 | ENSG00000132801.6 | 0.5 |
| ROMO1 | ENSG00000125995.15 | 0.5 |
| EPDR1 | ENSG00000086289.11 | 0.5 |
| DPM1 | ENSG00000000419.12 | 0.5 |
| JADE3 | ENSG00000102221.13 | 0.5 |
| PRPS2 | ENSG00000101911.12 | 0.49 |
| GS1-594A7.3 | ENSG00000225833.1 | 0.49 |
| RP11-243A14.1 | ENSG00000261617.1 | 0.49 |
| AHCY | ENSG00000101444.12 | 0.49 |
| AURKA | ENSG00000087586.17 | 0.49 |
| SLC5A6 | ENSG00000138074.14 | 0.49 |
| SCARB1 | ENSG00000073060.15 | 0.49 |
| SATB2 | ENSG00000119042.16 | 0.49 |
| SYAP1 | ENSG00000169895.5 | 0.49 |
| RALY | ENSG00000125970.11 | 0.49 |
| AGT | ENSG00000135744.7 | 0.49 |
| HSD17B10 | ENSG00000072506.12 | 0.49 |
| MAP7D2 | ENSG00000184368.15 | 0.49 |
| SLC19A3 | ENSG00000135917.13 | 0.48 |
| MFAP3L | ENSG00000198948.11 | 0.48 |
| TAF4 | ENSG00000130699.16 | 0.48 |
| DDX27 | ENSG00000124228.14 | 0.48 |
| TMLHE | ENSG00000185973.10 | 0.48 |
| PPP1R14D | ENSG00000166143.9 | 0.48 |
| DGAT2 | ENSG00000062282.14 | 0.48 |
| DBI | ENSG00000155368.16 | 0.48 |
| PLEKHG6 | ENSG00000008323.15 | 0.48 |
| REPS2 | ENSG00000169891.17 | 0.48 |
| TIMM17B | ENSG00000126768.12 | 0.48 |
| PGRMC1 | ENSG00000101856.9 | 0.47 |
| ARSE | ENSG00000157399.14 | 0.47 |
| YTHDF1 | ENSG00000149658.17 | 0.47 |
| MAPRE1 | ENSG00000101367.8 | 0.47 |
| FITM2 | ENSG00000197296.5 | 0.47 |
| SLC23A1 | ENSG00000170482.16 | 0.47 |
| PDRG1 | ENSG00000088356.5 | 0.47 |
| AAR2 | ENSG00000131043.11 | 0.47 |
| UQCC1 | ENSG00000101019.21 | 0.47 |
| COIL | ENSG00000121058.4 | 0.47 |
| YAE1D1 | ENSG00000241127.7 | 0.47 |
| TXLNG | ENSG00000086712.12 | 0.47 |
| PDHA1 | ENSG00000131828.13 | 0.47 |
| TSPAN33 | ENSG00000158457.5 | 0.47 |
| MYC | ENSG00000136997.14 | 0.47 |
| EIF6 | ENSG00000242372.6 | 0.47 |
| MTG2 | ENSG00000101181.17 | 0.47 |
| FAH | ENSG00000103876.11 | 0.47 |
| SHROOM4 | ENSG00000158352.15 | 0.47 |
| PIGU | ENSG00000101464.10 | 0.47 |
| TPX2 | ENSG00000088325.15 | 0.47 |
| KIF3B | ENSG00000101350.7 | 0.47 |
| EDA | ENSG00000158813.17 | 0.46 |
| TTPA | ENSG00000137561.4 | 0.46 |
| VAPB | ENSG00000124164.15 | 0.46 |
| CTD-2540B15.11 | ENSG00000267580.1 | 0.46 |
| SHROOM2 | ENSG00000146950.12 | 0.46 |
| PUS10 | ENSG00000162927.13 | 0.46 |
| COMMD7 | ENSG00000149600.11 | 0.46 |
| CTNNBL1 | ENSG00000132792.18 | 0.46 |
| LINC01555 | ENSG00000180869.4 | 0.46 |
| GPC4 | ENSG00000076716.8 | 0.46 |
| DDX52 | ENSG00000278053.4 | 0.46 |
| TSPY26P | ENSG00000235217.6 | 0.46 |
| PVR | ENSG00000073008.14 | 0.46 |
| A1CF | ENSG00000148584.14 | 0.46 |
| ASB9 | ENSG00000102048.15 | 0.46 |
| CHMP4B | ENSG00000101421.3 | 0.46 |
| BRCC3 | ENSG00000185515.14 | 0.46 |
| APEX2 | ENSG00000169188.4 | 0.46 |
| ADRM1 | ENSG00000130706.12 | 0.46 |
| UBE2V1 | ENSG00000244687.11 | 0.45 |
| SRC | ENSG00000197122.11 | 0.45 |
| PCTP | ENSG00000141179.13 | 0.45 |
| PHF20 | ENSG00000025293.15 | 0.45 |

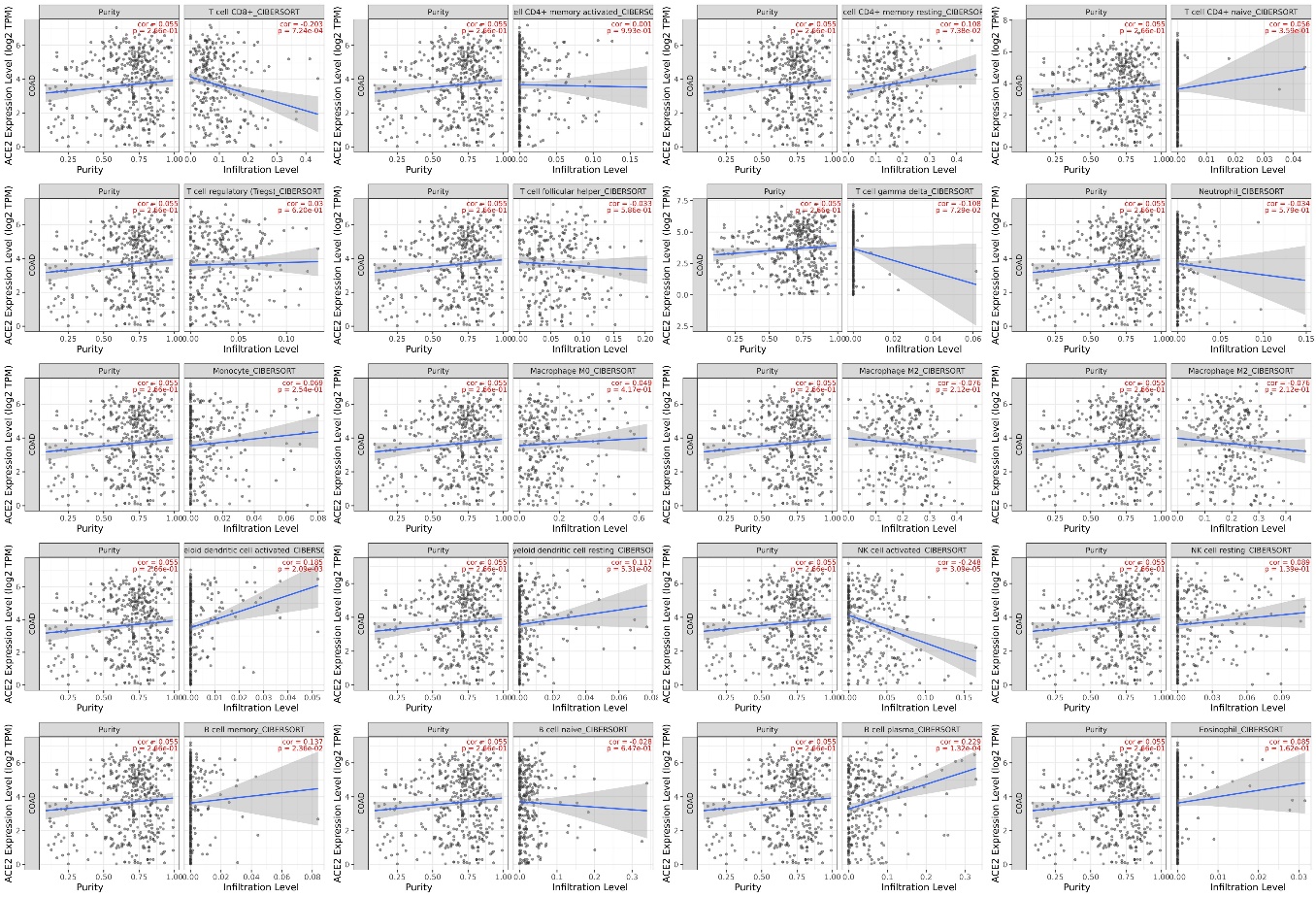

Figure S1. Correlation of the ACE2 gene expression with immune cell infiltration level in COAD. (CIBERSORT algorithm of TIMER 2.0 database)**.** The ACE2 was positively correlated with Memory B cell, Plasma B cell, activated myeloid DC, and resting myeloid DC, and negatively correlated with T cell, and activated NK cells. Colon Adenocarcinoma (COAD)
